# Towards a new treatment against polymicrobial infections: high antibacterial activity of lemon IntegroPectin against *Pseudomonas aeruginosa* and *Escherichia coli*

**DOI:** 10.1101/2020.06.16.154229

**Authors:** Alessandro Presentato, Antonino Scurria, Lorenzo Albanese, Pasquale Picone, Mario Pagliaro, Federica Zabini, Francesco Meneguzzo, Rosa Alduina, Domenico Nuzzo, Rosaria Ciriminna

## Abstract

Lemon IntegroPectin obtained via hydrodynamic cavitation of waste lemon peel in water only shows high antibacterial activity against two Gram-negative bacteria, *Pseudomonas aeruginosa* and *Escherichia coli*. The antibacterial effect against the ubiquitous pathogen *P. aeruginosa* was evaluated in terms of the minimal bactericidal (MBC) and minimal inhibitory concentration (MIC). Preliminary insight on the antibacterial mechanism of IntegroPectin originates from investigating its inhibitory activity against *E. coli*. Given the non-cytotoxic nature of citrus IntegroPectin and the ease of its reproducible production in large amounts, the route is open to the industrial development of a new antimicrobial treatment against polymicrobial infections unlikely to develop drug resistance.

## 1. Introduction

*Pseudomonas aeruginosa* is a Gram-negative bacterial strain ubiquitous in the environment due to its metabolic versatility and ability to degrade complex xenobiotics of anthropogenic origin.^[1]^ Unfortunately, this strain is also an opportunistic bacterial pathogen able to induce serious infections threatening human health.^[2]^ *P. aeruginosa*, for example, is one of the four frequently encountered bacteria responsible for causing hospital-acquired pneumonia, due to bacterial biofilm formation on endotracheal tubes in intubated patients.

Besides causing frequent infections of surgical sites, and of chronic decubitus ulcers, *P. aeruginosa* is also involved in infections of the urinary tract,^[3]^ and in cystic fibrosis promoting accelerated decline of pulmonary function. Unfortunately, the bacterium exhibits significant resistance to both innate immune anti-microbial peptides and to several antibiotics.^[4]^

The increase of antibiotic resistant phenotypes and antibiotic resistance genes in the environment^[5]^ and in widely different organisms,^[6]^ requires to urgently identify and develop at industrial level new antimicrobial solutions against said pathogens.

To identify the targets for future antibacterials capable of preventing biofilm formation on indwelling plastic tubes frequently used in clinical settings, scholars in India recently identified the 56 specific proteins in highly virulent *P. aeruginosa* PA14 strain regulating its ability to form biofilm on said tubes.^[7]^

In general, to manage *Pseudomonas aeruginosa* infections, causing high mortality in critically ill and immunocompromised patients driven by the appearance of drug-resistant strains, today’s therapeutic options include antibiotic combinations based on pharmacokinetic and pharmacodynamic analyses.^[8]^

Highly desirable new anti-pseudomonal medicines should not drive superinfection,^[8]^ and be preferably available as oral formulations to allow step-down therapy in the treatment of Gram-negative bloodstream infection.^[9]^

Similarly, *Escherichia coli* is a versatile Gram-negative bacterial strain colonizing the human gastrointestinal tract, where it becomes part of the microbiota a few hours after birth. Specific pathotypes of *E. coli* may infect healthy individuals affording three possible syndromes: enteric disease and diarrhea, urinary tract infection, and meningitis.^[10]^

In a previous study, lemon IntegroPectin was shown to exert significant activity *in vitro* against *Staphylococcus aureus* virulent strains.^[11]^ We now report the discovery of high *in vitro* activity of lemon IntegroPectin against virulent strains of *P. aeruginosa*, evaluating its antibacterial effect in terms of the minimal bactericidal (MBC) and minimal inhibitory concentration (MIC). We also show that this new biomaterial inhibits *E. coli* proliferation by exerting a powerful oxidative stress action against the microorganism.

## 2. Results and Discussion

Lemon IntegroPectin (IntPec) is effectively and reproducibly produced via hydrodynamic cavitation of waste lemon peel in water only. The lemon peel was derived by an in-line extractor at a lemon juice factory and directly processed on semi-industrial scale (34 kg of waste lemon peel of organically grown Siracusa lemons in 120 L tap water) using the same cavitation conditions lately developed to process waste orange peel.^[12]^

Easily dissolved in water, the resulting yellow pectin flakes obtained after freeze drying have a pleasant lemon smell and are completely devoid of cytotoxic activity against human pulmonary cells up to high concentration (1 mg/mL).^[13]^ Furthermore, lemon IntPec has an extremely high content of polyphenols adsorbed at its surface: 0.88 mg GAE/g (in terms of gallic acid equivalents, or GAE, per dry gram of pectin) versus 8.3×10^−3^ mg GAE/g for the lemon peel of the cultivar with the highest biophenol concentration.

We tested the antimicrobial activities of IntPec against *P. aeruginosa* 10145 strain (indicator pathogen strain) commonly used as a positive control for molecular detection in bioaerosols, as well as a quality control strain for drugs.^[14]^ For comparison, we also tested the antibacterial activity of commercial citrus pectin (galacturonic acid ≥74.0%, dry basis, from Merck Life Science, Milan, Italy).

Figure 1 shows evidence that both commercial pectin and IntPec inhibit *P. aeruginosa* growth. However, the number of viable cells decreased by 2.6 log units when the concentration of IntPec went from 0 (control) to 10 mg mL^-1^, whereas it remained almost unvaried (from 9.1 to 9.0) when the concentration of commercial citrus pectin was 10 mg mL^-1^.

**Figure 1.**
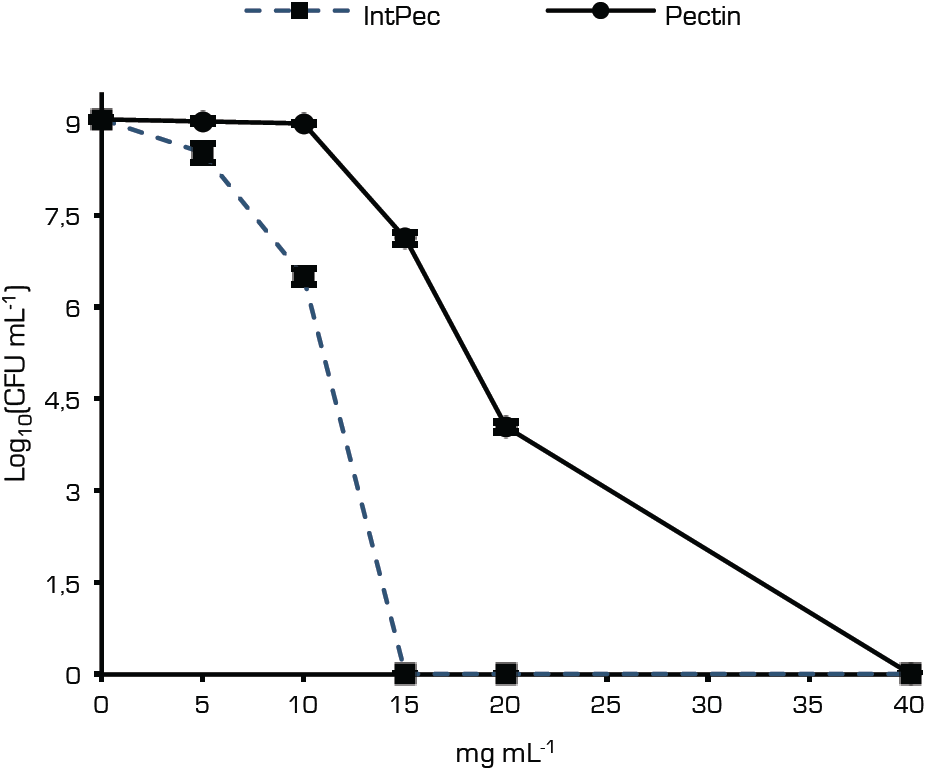
Viable cell count of *P. aeruginosa* 10145 strain after incubation in culture broth added with increasing concentrations of IntPec and commercial citrus pectin.

Increasing the concentration of IntegroPectin from 10 to just 15 mg mL^-1^, the Log_10_ (CFU) reached the 0 value, obtaining a total killing effect and establishing the MBC of the lemon IntegroPectin against *P. aeruginosa*. At the same concentration of commercial citrus pectin, the Log_10_ (CFU) was still 7.1. A concentration of 40 mg mL^-1^ of commercial pectin was required to observe Log_10_ (CFU) to reach the 0 value.

Evidence of the reduction of viable *P. aeruginosa* 10145 CFUs challenged *in vitro* with pectin and IntPec is displayed in Figure 2.

**Figure 2.**
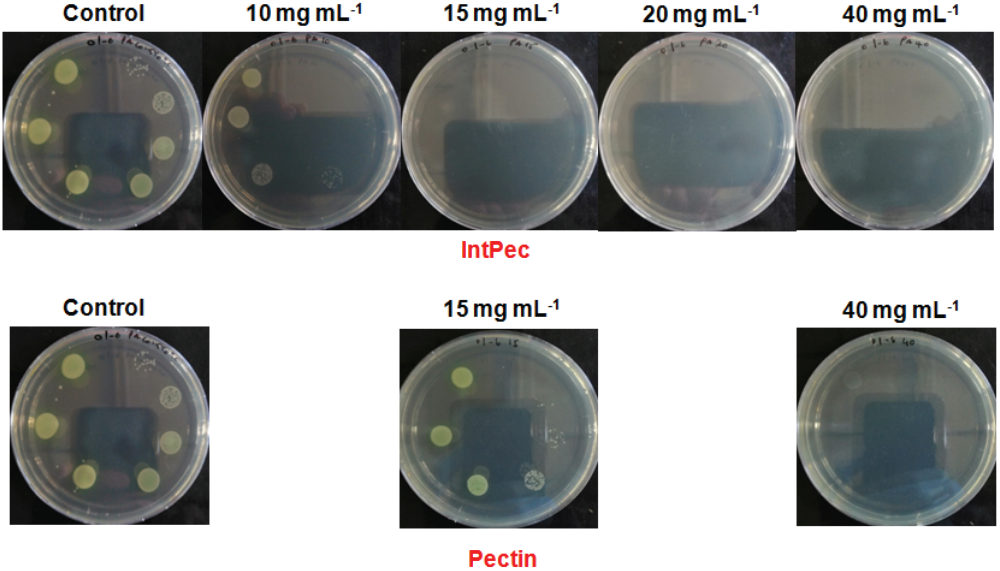
Evidence of *in vitro* pectin and IntegroPectin activity against *P. aeruginosa* 10145 strains.

To establish MIC values of both pectins, we measured the optical density at 600 nm (at this wavelength the bacterial cells are not harmed whereas light scattered by the cells no longer reaches the photoelectric cell translating into higher turbidity) of the challenged *P. aeruginosa* cultures as compared to unchallenged ones. We briefly remind that the MIC value is defined as the lowest concentration of an antimicrobial agent that prevents visible growth of a microorganism in a broth dilution susceptibility test.^[15]^

Figure 3 shows that the MIC value of lemon IntPec is 10 mg mL^-1^ whereas that of commercial citrus pectin is 20 mg mL^-1^.

**Figure 3.**
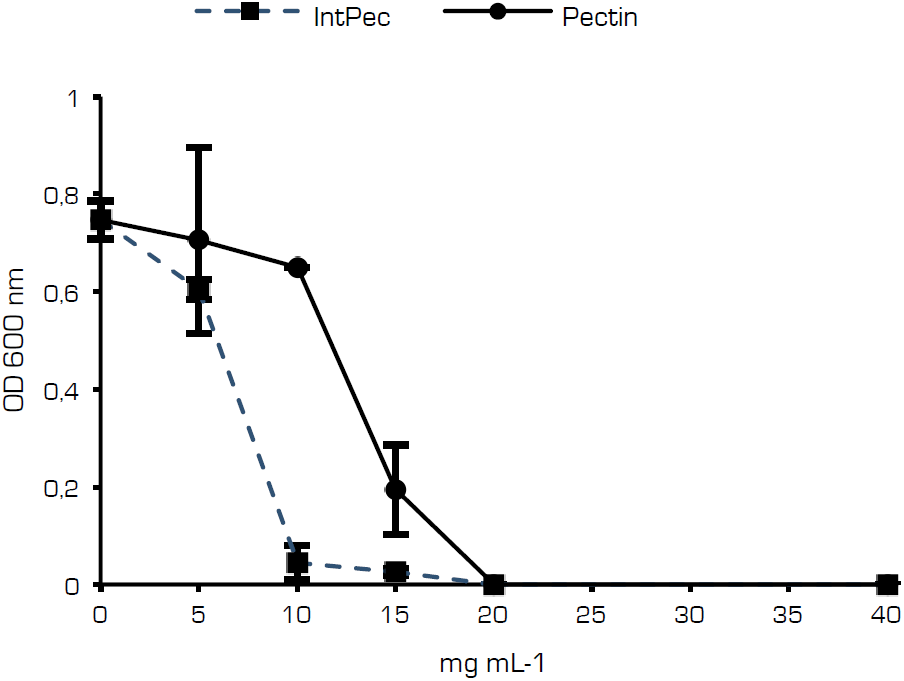
Optical density at 660 nm of *P. aeruginosa* ATCC 10145 cultures in the presence of lemon IntPec and of commercial citrus pectin for the assessment of minimal inhibitory concentration (MIC).

To better understand the relevance of these results, it is instructive to realize that the recommended broth dilution method to determine the minimal inhibitory concentration of antimicrobial substances consists of a 5×10^5^ CFU mL^-1^ inoculum size,^[16]^ while in the present study on the efficacy of lemon IntPec against *P. aeruginosa*, the broth was inoculated with a cellular load 2 orders of magnitude higher than the recommended bacterial concentration.

To provide preliminary insight on the antibacterial mechanism, the effect of IntPec was also tested on *Escherichia coli* growth by incubating cells in presence of sub-inhibitory concentrations of IntPec.

Figure 4 shows evidence of the increasing antibacterial activity of IntPec at increasing concentrations.

**Figure 4.**
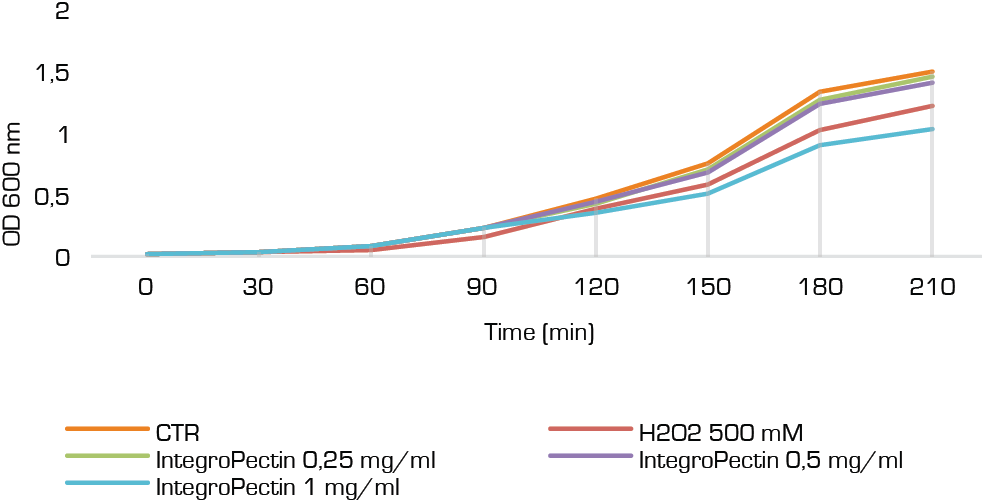
Effect of H_2_O_2_ 500 mM and of lemon IntPec increasing concentrations on *E. Coli* proliferation. CTR stands for control sample.

While little activity is observed at 0.25 mg/mL titre, it is enough to add the lemon IntPec at 1.0 mg/mL concentration to observe a dramatic reduction in the *E. coli* cell proliferation, significantly higher than that caused by concentrated (0.5 M) aqueous hydrogen peroxide (H_2_O_2_), a powerful oxidizing agent that at this concentration is capable to quickly denature enzymes and oxidize not only protein side chains but also the protein backbone.^[17]^

To evaluate the possible generation of Reactive Oxygen Species (ROS), *E. coli* bacteria were incubated with a diluted solution of dichlorodihydrofluorescein diacetate (DCFH-DA), the most widely used probe for detecting intracellular H_2_O_2_ and oxidative stress,^[18]^ alone or in the presence of increasing concentrations of IntPec.

Figure 5 shows that already at 0.5 mg/mL concentration the bioproduct exerts a powerful oxidative stress on the bacteria, significantly higher than H_2_O_2_ 0.5 M. When the IntPec concentration is increased to 1.0 mg/mL, the ROS generation becomes more than twice higher than that driven by concentrated H_2_O_2_.

**Figure 5.**
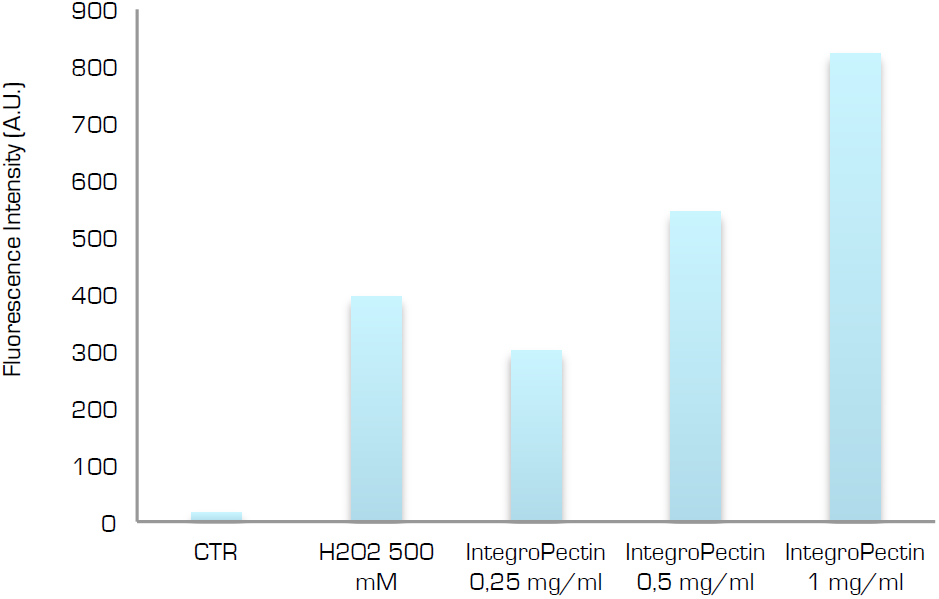
Generation of ROS on *E. coli* bacteria driven by aqueous H_2_O_2_ 0.5 M and by increasing concentrations of lemon IntPec. CTR stands for control sample.

Figure 6 shows the quick production of ROS estimated again using DCFH-DA after increasing the IntPec concentration from 0.25 mg/mL to 1 mg/mL. The amount of ROS formed at the surface of *E. coli* follows an almost identical profile for H_2_O_2_ 500 mM and aqueous IntPec at 0.5 mg/mL concentration.

**Figure 6.**
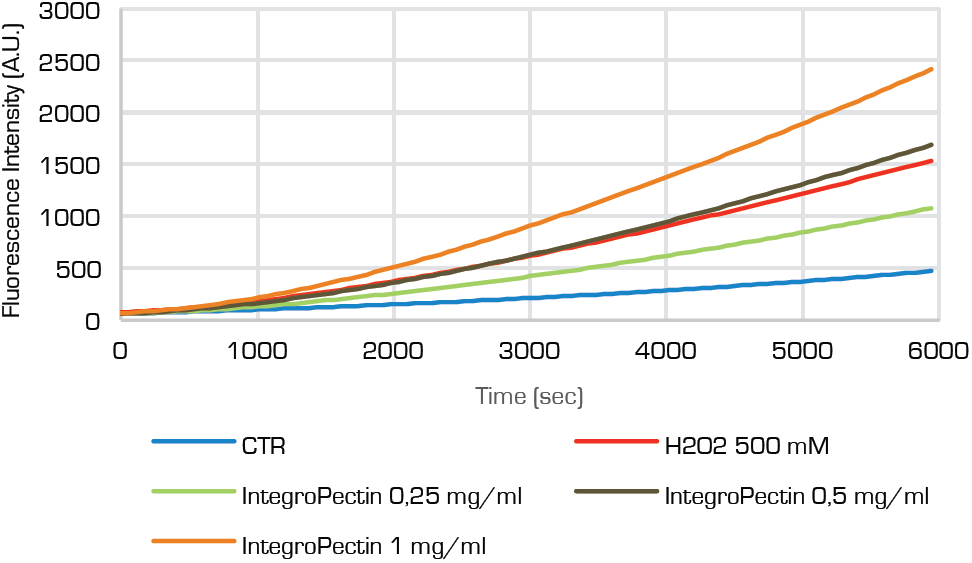
*E. coli* oxidation kinetics driven by by aqueous H_2_O_2_ 0.5 M and by increasing concentrations of lemon IntPec. CTR stands for control sample.

Yet, using lemon IntPec at 1.0 mg/mL already after 16 min (∼1000 s) the amount of ROS is higher than that generated by concentrated hydrogen peroxide, to eventually almost double the oxidative stress exerted by H_2_O_2_ 0.5 M after 1.7 h since the addition of the new pectic substance.

The same lemon IntPec was recently shown to exert significant activity *in vitro* against *S. aureus* virulent strains.^[11]^ It is therefore of direct relevance to this account the fact that Gram-positive *S. aureus* is often present in conjunction to *P. aeruginosa* forming hazardous polymicrobial complex communities particularly resistant to antibiotics.^[19]^

Originally reported in the late 1930s,^[20]^ the antibacterial properties of pectin were rediscovered in the late 1990s by scholars in Russia. Pectin, usually derived from citrus peel or apple pomace, was found to be the only food fiber showing bactericidal activity on the most widely distributed pathogenic and opportunistic microorganisms, decreasing the need for antimicrobial penicillins.^[21]^

In 2015, scholars in Lebanon reported the antibacterial activity of citrus pectin against *S. aureus*, with optimal antibacterial activity observed at pH 6 with MIC values ranging from 0.39 mg mL^-1^ to 3.125 mg mL^-1^.^[22]^ However, scholars in Taiwan in 2014 reported lack of any antibacterial activity of citrus pectin against *P. aeruginosa*.^[23]^

We make the hypothesis that the antibacterial action of lemon IntegroPectin is due to a synergistic combination of *i*) the pectic polymer, *ii*) the lemon oil adsorbed at its surface (the citrus essential oil is present in the form of a highly stable nanoemulsion in the aqueous phase after the hydrocavitation process),^[12]^ and *iii*) the lemon flavonoids highly concentrated (with respect to the citrus peel) at the surface of lyophilized IntPec.^[13]^

Likewise to other citrus oils, lemon oil is particularly active against Gram-positive bacteria, even though selected Gram-negative bacteria such as *E. coli* and *Campylobacter jejuni* are inhibited by lemon oil due to the presence of oxygenated monoterpenes exhibiting much stronger antimicrobial activity than hydrocarbon monoterpenes present in the essential oil.^[24]^

These results were subsequently confirmed for the fungicidal action of lemon oil against *Candida albicans* pathogenic yeasts, wherein the most active antimicrobial was found to be citral.^[25]^ Remarkably, the concentration of the latter oxygenated monoterpene is particularly abundant in the peel of citrus grown (in Sicily) according to organic farming principles.^[26]^ Indeed, the waste lemon peel used to produce the IntPec was obtained by a citrus company in Sicily using only organically grown lemons.

Finally, citrus flavonoids chiefly contained in the flavedo and in the albedo are well known antibacterials.^[27]^ Only recently the mechanism through which they inhibit *P. aeruginosa* biofilm formation was identified. Flavonoids possessing dihydroxyl moieties in the flavone A-ring backbone prevent LasR DNA binding due to the inhibitory action of the two hydroxyl groups, with one of them that must be at position 7 on the A-ring for potent inhibition of the aforementioned binding.^[28]^

In doing so, citrus flavonoids inhibit quorum sensing (QS, the bacterial cell-cell communication process controlling collective behavior) via antagonism of the QS receptor, the LasR protein. The latter protein indeed is a major transcriptional activator of *P. aeruginosa* QS and plays a pivotal role in the activation of many virulence genes.^[29]^

Administration of citrus flavonoids to *P. aeruginosa* alters transcription of quorum sensing-controlled target promoters and suppresses virulence factor production, confirming their potential as anti-infectives that do *not* function by means of traditional bactericidal or bacteriostatic mechanisms.^[28]^

All main lemon flavonoids, hesperidin (hesperetin 7-*O*-beta-rutinoside), eriocitrin (eriodictyol 7-*O*-beta-rutinoside), and diosmin (diosmetin 7-*O*-rutinoside) are the glycosides of flavonoids of similar compounds bearing two dihydroxyl moieties in the flavone A-ring backbone, with the hydroxyl at position 7 on the A-ring derivatised with a sugar.^[30]^ The latter glycosidic bond is easily hydrolyzed at the bacterial membrane level in the aqueous culture broth.

Accordingly, orange extracts rich in glycosylated flavanones including hesperidin, naringenin, and naringin are known to effectively interfere and inhibit quorum sensing in *P. aeruginosa*^[31]^ as well as in Gram-negative *Yersinia enterocolitica*^[32]^ bacteria.

## 3. Conclusions

In conclusion, we have discovered the powerful *in vitro* activity of lemon IntegroPectin against virulent strains of *P. aeruginosa* and *E. coli*. Aiming to identify the mechanism of action of this novel bioproduct, we have also discovered that dissolved in solution at the concentration of 1 mg mL^-1^, it exerts an oxidative stress action against the latter microorganism that is more than twice stronger than concentrated (0.5 M) hydrogen peroxide.

Taking into account the complete lack of cytotoxicity of citrus IntPec against human lung cells at the aforementioned concentration (1 mg mL^-1^),^[13]^ and the antibacterial activity of lemon IntPec against Gram-positive *S. aureus*,^[11]^ these results may open the route to the industrial development a new antibacterial treatment against polymicrobial infections based on a new bioproduct reproducibly and economically obtained in large amounts from the main citrus industry’s by-product.

### Experimental

#### Experiments with P. aeruginosa

The antibacterial activity of IntePec and commercial citrus pectin was preliminarily assessed through the determination of MBC and MIC. Specifically, stationary grown cells of *P. aeruginosa* were inoculated (1.19×10^7^±8.01×10^5^ CFUmL^-1^ *n* = 6) in Luria Bertani medium (hereafter named as LB and composed of [g L^-1^] sodium chloride [10], trypton [10], and yeast extract [5]) amended with increasing concentrations (i.e., 5, 10, 15, 20, and 40 mg mL^-1^) of either lemon IntPec or commercial citrus pectin, being then challenged for 24 h at 37°C under mechanical shacking (180 rpm).

Non-challenged bacterial cultures were incubated under the same conditions and used as a control. After 24 h challenge, all the bacterial cultures were serially diluted and aliquots (20 µL) of the diluted cultures, as well as the undiluted ones, were spotted onto LB agar (15 g L^-1^) plates and recovered at 37°C under static mode.^[33]^ The kill curve reporting the number of viable CFU mL^-1^ as a function of the concentration of both pectins is expressed in logarithmic (Log_10_) scale with standard deviation (*n* = 3), as described elsewhere.^[34]^

#### Experiments with E. coli

A single colony was inoculated into LB medium and incubated at 37°C overnight (o.n.). An aliquot (5 µL) of the o.n. bacterial culture, approximately 10^9^ CFU/mL, was added to three test tubes containing fresh LB medium (5 mL). IntPec (0.25, 0.5 and 1 mg/mL) was added separately to the culture medium at time 0 min or 210 min. The growth was determined by reading the absorbance value at 600 nm (OD600) using a Spark 10 M multimode microplate reader (Tecan Group, Männedorf, Switzerland) with 30 min intervals.

For ROS generation, an aliquot of the *E. coli* o.n. culture, approximately 10^9^ CFU/mL, was diluted (1:10^5^) and a 100 µL sample placed in a 96-well optical bottom white microplate. IntPec at different concentration (0.25, 0.5 and 1 mg/mL) was added to the wells. Then, the samples were incubated with 1 mM of DCFH-DA (Molecular Probes, Eugene, OR, USA) for 8 h at 37°C. Afterward, the *E. coli* samples were analyzed by using a GloMax Microplate Reader (Promega, Madison, WI, USA) for fluorescence detection at the excitation wavelength of 475 nm and emission wavelength 555 nm. An untreated *E. coli* bacterial culture (CTR) and an *E. coli* culture challenged with 500 mM H_2_O_2_ were used as control for growth and oxidative stress.

For oxidation kinetics, an aliquot of *E. coli* o.n. culture, approximately 10^9^ CFU/mL, was diluted (1:10^5^) and a 100 µL sample placed in a 96-well optical bottom white microplate. IntPec at different concentration (0.25, 0.5 and 1 mg/mL) was added to the wells. The production of ROS was estimated using DCFH-DA at a final concentration of 1 mM. Untreated *E. coli* (CTR) was used as control and aqueous H_2_O_2_ 0.5 M was used as a positive control of oxidation state. The oxidation kinetics was followed by fluorescence at the excitation wavelength of 475 nm and emission wavelength 555 nm by a GloMax Discover System (Promega, Madison, WI, USA) in a 96-multiwell plate incubated for 2 h at 37°C.

## Acknowledgements

Thanks to Campisi Italia (Siracusa, Italy) for the gift of waste lemon peel from which the IntegroPectin used in this study was obtained. We thank Dr M. Di Carlo, CNR-IRIB, for purposeful collaboration.

## Author Information

### Notes

The Authors declare no conflict of interest

